# *In Vitro* Potent Activity of ε-poly-L-lysine against *Candida albicans* and the Underlying Mechanisms

**DOI:** 10.1101/605857

**Authors:** Lian-hua Wei, Tian Yu, Xiao-ning Wang, Jin-xia Hou, Xin Wang, Chuan Wang, Ke-ke Li, Shuang-yan Jing, Xu Zhang

## Abstract

**Objective:** This study aimed to examine the antifungal activity of ε-poly-L-lysine (ε-PL) against the planktonic cells or biofilms of *Candida albicans* and explore the underlying mechanism.

**Methods:** The minimal inhibitory concentration, minimum fungal concentration, and sessile minimal inhibitory concentration were estimated. The germ tube formation and yeast-to-hypha transformation of *C. albicans* in different media that induced mycelial growth were recorded. The effect of different concentrations of ε-PL on the biofilm formation process and mature biofilm of *C. albicans* was determined. The reactive oxygen species (ROS) and malondialdehyde (MDA) contents of *C. albicans* after ε-PL treatment were measured. The changes in major virulence genes and proteins of *C. albicans* were detected.

**Results:** ε-PL (512 μg/mL) exerted a strong inhibitory effect on *C. albicans* and biofilms. It blocked the yeast-to-hypha transition and reduced the germ tube formation and germ tube length of *C. albicans*. The MDA and ROS contents showed an upward trend, indicating a positive correlation with the concentration. Further, ε-PL inhibited the high expression of virulence genes in oxidative stress induced by *C. albicans*. The main peak in the mass spectrum of *C. albicans* was found to be clear.

**Conclusions:** ε-PL exerted a significant antifungal effect on the phytoplankton and biofilm of *C. albicans*. High concentrations of ε-PL significantly inhibited the main mycelium of *C. albicans*. ε-PL induced ROS, released cytochrome C, attacked the *C. albicans* cell membrane to aggravate its lipid oxidation, and inhibited the expression of *C. albicans*–associated virulence genes and proteins, thereby exerting a bacteriostatic effect.

**Importance:** The last two decades have seen a growing trend toward the failure of current antifungal drugs attributed to *Candida* biofilms. Under appropriate conditions, adherence and colonization of planktonic cells on host tissues and medical devices initiate multicellular organization called biofilm, which is an organized heterogeneous mixture of yeast, hyphae, and pseudohyphal forms embedded in a complex extracellular matrix. Compared with the planktonic cells, biofilms show high resistance to a wide variety of antifungal agents and tolerance to harsh environments and host immune system. Moreover, the development of antifungal drugs is costly, long-term, and difficult. Thus, researchers turned their attention to natural antibacterial peptides, hoping to find an effective antifungal substance or enhance the sensitivity of the existing antifungal drugs to *C. albicans*.

## Introduction

*Candida albicans*, also known as *Canadian albicans*, is widely found in nature, and is often parasitic on human skin, mucous membranes, oral cavity, upper respiratory tract, and vagina^[1-2]^. Nowadays, with the increase in the number of immunocompromised patients and patients with medically implanted devices (catheters, heart valves, cardiac pacemakers, vascular bypass grafts, endotracheal tubes, and central nervous system shunts), fungal infection has become an important issue that cannot be ignored in clinical practice, and *Candida* disease, represented by *C. albicans*, is a hot issue^[3]^. According to statistics, *C. albicans* is the fourth largest pathogen of systemic infection in the United States, with a mortality rate of up to 50% ^[4]^. Although people are trying to curb this situation, due to delayed diagnosis and antifungal resistance, candidiasis is associated with high mortality worldwide. The last two decades have seen a growing trend toward the failure of current antifungal drugs attributed to *Candida* biofilms ^[5-6]^. Under appropriate conditions, adherence and colonization of planktonic cells on host tissues and medical devices initiate multicellular organization called biofilm, which is an organized heterogeneous mixture of yeast, hyphae, and pseudohyphal forms embedded in a complex extracellular matrix ^[7-8]^. Compared with the planktonic cells, biofilms show high resistance to a wide variety of antifungal agents and tolerance to harsh environments and host immune system ^[9]^. Moreover, the development of antifungal drugs is costly, long-term, and difficult. Thus, researchers turned their attention to natural antibacterial peptides, hoping to find an effective antifungal substance or enhance the sensitivity of the existing antifungal drugs to *C. albicans* ^[10]^.

Antibacterial peptides, a class of natural antimicrobial substances, show anti-adhesive and antimicrobial activities. Particularly, ε-poly-L-lysine (ε-PL) has a broad antibacterial spectrum, high heat resistance, and good antibacterial action, even in acidic or alkaline medium. Especially, it has bacteriostatic effects on Gram-negative bacteria, such as *Escherichia coli* and *Salmonella*, which are not inhibited by other natural preservatives (such as nisin and natamycin)^[11]^. As early as 2003, Food and Drug Administration(FDA)approved it as a new type of natural food preservative, and related research focused on its antibacterial properties and production and purification techniques as preservatives. Only a few publications mentioned that ε-PL could strip the outer membrane and cause an abnormal distribution of the cytoplasm ^[12]^. Although some studies have been carried out on the antifungal mechanism of ε-PL, none exhibited the specific molecular mechanism of ε-PL on fungi^[13]^. To promote the application of ε-PL, elucidating its antibacterial mechanism is particularly important. This study aimed to confirm the antibacterial activity of ε-PL against *C. albicans* and its inhibitory effect on *C. albicans* biofilms. At the same time, its antibacterial mechanism was initially explored for its clinical application, thus providing a theoretical basis for new ideas and directions for clinical antifungal drug research and development.

## Methods

### Strain and growth conditions

The standard strains of *C. albicans*, ATCC 64548 and ATCC 64550, were purchased from American type cell culture (ATCC). Clinical isolates (CA1-50) were purified from vaginal, fecal, sputum, and lavage fluid samples, which were identified by Matrix-assisted Laser Desorption-ionization Time of Flight Mass Spectrometry (MALD-TPF-MS). All strains were routinely grown in Sabouraud Dextrose Agar (SDA) at 30°C in a shaking incubator.

### Antifungal susceptibility *in vitro*

The minimal inhibitory concentrations (MIC) of ε-PL against all the 52 strains were determined according to the broth microdilution protocol of Clinical and Laboratory Standards Institute(CLSI)with minor modifications ^[14]^. Briefly, the initial fungal suspension used for inoculation was adjusted to 10^3^ CFU/mL in the Roswell Park Memorial Institute-1640 medium and the final concentrations of ε-PL were adjusted from 16 to 4096 μg/mL. The plates were incubated at 37°C for 48 h. Growth inhibition was determined using optical density at 490 nm (OD_490_), from which the background ODs were already subtracted. The MIC_80_ was defined as the concentration of drugs that inhibited 80% of the cell growth. Each strain was tested in triplicate with positive and negative growths and drug controls.

### Antibiofilm susceptibility *in vitro*

The antibiofilm effect of ε-PL was measured using crystal violet (CV) as described in a study by Feng with minor modifications ^[15]^. Briefly, 96-well plates were seeded with 200-μL aliquots of 2 × 10^3^ CFU/mL fungal suspension and incubated statically at 37°C for 1.5 h for initial adhesion. Then, the 96-well plates were washed slightly with phosphate-buffered saline (PBS; pH 7.2) to remove nonadherent cells. A fresh RPMI-1640 medium with different concentrations of ε-PL was added, followed by another 24 h of incubation. Then, the cells were crystallized after dry fixation for 3 h, followed by rinsing with flowing distilled water until the medium was colorless. The cells were then dried for 2 h, and 150 μL of 30% glacial acetic acid was added to each well. Growth inhibition was determined using a spectrophotometer. OD_590_ was measured, and the sessile minimal inhibitory concentration (SMIC_50_) was defined as the concentration of drugs that inhibited 50% of cell growth.

### Time-kill curve assay

For the time-kill curve assay, the *C. albicans* cells (ATCC64548 and ATCC64550) in the exponential growth phase were resuspended in saline and adjusted to 5 × 10^4^ CFU/mL. Different concentrations of ε-PL formulated with the RPMI-1640 medium were added to the suspensions. The samples were cultured at 37°C under constant shaking (200 rpm), and small aliquots were withdrawn at predetermined time points (0, 3, 6, 9, and 12 h). The experiments were performed in triplicate ^[16]^.

### *In vitro* antibiofilm assay

To determine the *in vitro* assay of ε-PL against mature biofilms, *C. albicans* cells (ATCC64548 and ATCC64550) grown overnight in the Yeast Extract Peptone Dextrose Medium(YPD)medium were centrifuged and washed in PBS. A suspension of 1 × 10^6^ cells/mL was prepared in physiological saline and inoculated into 96-well plates. Oral plates were inoculated with 100 μL of 1 × 10 ^7^ CFU/well. The bacterial suspension was incubated for 90 min at 37°C and 75 rpm. After adhering of the fungi, the suspension was discarded and washed twice with PBS. The supernatant was aspirated to remove unattached cells. A fresh RPMI-1640 medium with or without different concentrations of ε-PL was added and incubated for an additional 24 h. To detect ε-PL on mature biofilms, 100 μL of the RPMI-1640 medium and 100 μL of different concentrations of ε-PL were added. The concentration was 0, 1/2 MIC, MIC, and Minimum Fungs concentration (MFC), with shaking at 37°C and 75 rpm for 24 h. The degree of clarity of ε-PL against mature biofilms of *C. albicans* was determined using the SMIC_50_ method [17].

To determine the effect of ε-PL on biofilm formation, biofilm biomass was indirectly measured using CV staining (70). After incubating the preserved cells at 37°C for 90 min, the biofilm was washed three times with sterile PBS to remove unattached cells. ε-PL was diluted to 11 different concentrations and inoculated into the treated 96-well plates at working concentrations of 16, 32, 64, 128, 256, 512, 1024, 2048, and 4096 μg/mL. Then, a blank control group was set. After sealing with a plastic paper and placing it in a fungal growth incubator at 35°C for 24 h, the biofilm formation was measured using the CV method. The experiment was performed in triplicate, and the mean absorbance of each well was plotted against the ε-PL concentration.

### Confocal laser scanning microscopy

The *C. albicans* ATCC64548 was cultured on the SDA medium overnight and then inoculated in the YPD liquid medium at 35°C and 200 rpm. The culture was shaken to log phase, and after collecting the cells, the physiological saline was used to prepare suspensions of 2 × 10^6^ CFU/mL. The prepared bacterial suspension was added to a laser confocal culture dish for 37 min at 37°C and 200 rpm and then washed two to three times with PBS. Then, 1 mL of RPMI-1640 liquid medium and 1 mL of different concentrations of ε-PL were added. Suspensions with different working concentrations of ε-PL (0, 256, 512, and 1024 μg/mL) were cultured at 37°C for 24 h. Then, the cells were co-stained with 10 μg/mL fluorescein diacetate (FDA) and 10 μg/mL propidium iodide (PI) at 37°C for 45 min in the dark. Images were taken using a confocal laser scanning microscope (CLSM).

### Filamentation assay

The activated *C. albicans* ATCC64548 was formulated into 2 × 10^5^ CFU/mL using physiological saline and then inoculated separately in the RPMI-1640, YPD liquid containing 10% serum, and Spider media. Different concentrations of ε-PL were added to the bovine serum to make working concentrations of 0, 1/4 MIC, 1/2 MIC, MIC, and MFC at 37°C for 3, 6, and 9 h. The morphology of the mycelium was detected using an inverted microscope. Each experiment was repeated three times.

### Measurement of intracellular reactive oxygen species

The intracellular reactive oxygen species (ROS) in *C. albicans* ATCC64548 biofilm cells was measured using 2,7-Dichlorodi -hydrofluorescein diacetate (Sigma, the US) as described in a previous study ^[19]^. Briefly, the ε-PL-treated cells were pooled by centrifugation (3000 g, 5 min), resuspended in sterile PBS to a final concentration of 6 × 10^6^ CFU/mL, stained with DCFH-DA in the dark at 37°C for 30 min, and finally analyzed using a fluorescence spectrophotometer (F-4500, HITACHI Instrument, Tokyo, Japan).

### Determination of lipid oxidation by treating *C. albicans* with ε-PL

To activate *C. albicans* ATCC64548, physiological saline configured to a concentration of 0.5 McMillan suspension was used. The strains were divided into groups A and B. Group A treatment method was as described earlier. For group B, 10mM N-acetylcysteine (NAC) was added in group A. The strains in the two groups were allowed to vortex at 37°C and 200 rpm for 12 h^[20]^. Then they were centrifuged at 5000*g* for 5 min to collect the bacteria. Total protein was extracted using the Yeast Protein Extraction Kit(Omega, China), and the protein concentration was measured using the BCA Protein Assay Kit(Sangon Biotech, China). The lipid oxidation (malondialdehyde, MDA) test kit (Wanleibio, China)was used to determine lipid oxidation damage. Each set of experiments was repeated three times.

### Effect of ε-PL on the cell membrane of *C. albicans* detected by PI staining

After preserving the standard strains ATCC 64548 (fluconazole-sensitive strain) and ATCC 64550 (fluconazole-resistant strain) of *C. albicans* in the SDA medium, they were configured to 1 × 10^7^ CFU/mL using physiological saline and diluted 200 times with an RPMI-1640 medium. Then, different concentrations of ε-PL were added to 500 μL of diluted bacteria solution, and, and the working concentration was adjusted to 0, 256, 512, and 1024 μg/mL. After incubating for 12 h at 37°C, the cells were collected at 1000 rpm and washed two to three times with PBS buffer. They were added to 250 μL of PBS and stained with 5 μg/mL PI at 30°C. The cells were then incubated in a water bath for 2 h, centrifuged at 3000 rpm for 5 min, washed three times with sterile PBS, and observed using a fluorescence microscope.

### Quantification of gene expression by real-time reverse transcriptase–polymerase chain reaction

Real-time reverse transcriptase–polymerase chain reaction (RT-PCR) was conducted for validating the results derived from the RNA-seq (31). Total RNAs were isolated from the preformed biofilms using the hot-phenol method. Approximately 1 g of total RNA was used to synthesize cDNA using a ReverTra Ace quantitative PCR RT Kit (Toyobo Ltd., Osaka, Japan). Then, cDNA was diluted (1:5), followed by amplification with Synergy Brands real-time PCR master mix (Toyobo Ltd.) in a final volume of 20:l. Quantitative RT-PCR was carried out using a LightCycler 480 instrument (Roche Molecular Biochemicals, Mannheim, Germany). The relative differences in mRNA abundance thus obtained were calculated by determining the changes in the cycle threshold values. The data were normalized to the level of expression of *act* as its expression level remained unchanged in both yeast cells and biofilms. The relative fold change in the gene expression levels was calculated using the 2ΔΔCT method.

### Determination of changes in the protein profile of *C. albicans* induced by ε-PL

Baicalin was added to *C. albicans* ATCC64548 biofilm cell suspension (2 × 10^6^ CFU/mL) to the final concentrations of 0, 256, 512, and 1024 μg/mL of ε-PL and cultured for 12 h with shaking. After centrifuging the suspension and leaching it twice with PBS, 5–10 mg was added to a 1.5-mL centrifuge tube containing 300 μL of sterile water and mixed well. Then, 900 μL of ethanol was added, vortexed, and mixed. The mixture was centrifuged at 12,000 rpm for 3 min. The supernatant was discarded and centrifuged at 12,000 rpm for 1 min. A pipette was used to aspirate the supernatant, and the residual liquid was allowed to evaporate. Then, 50 μL of extract A was added to the cell pellet, mixed with a pipette, and allowed to stand for 2 min at room temperature. Glass beads were added and ground with a rod for 3 min. Extract B was added, vortexed and mixed, and centrifuged at 12,000 rpm for 3 min. Then 1 μL of the supernatant was taken in a pipette and added to the sample of the target plate at room temperature and allowed to dry. Next, 1 μL of the matrix solution was added to cover the sample, allowed to dry naturally, and tested on the machine. The procedure was repeated 10 times. Mass spectrometry analysis was used to map different groups of samples.

## Results

### Antifungal activity of ε-PL

Supplement Table 1 shows that ε-PL had a broad-spectrum antifungal ability, exerting a potent to moderate killing effect, especially against the fluconazole-resistant strain.

The MIC_80_ was between 256 and 512 μg/mL against clinical isolates and standard strains (the fluconazole-sensitive strain ATCC64548 and the fluconazole-resistant strain ATCC64550). The time-kill assay showed that ε-PL inhibited the growth of *C. albicans* SC5314 cells in a dose-dependent manner (Fig. 1). A striking fact about the figures is that MIC and MFC resulted in a more than 100-fold reduction in the numbers of viable cells compared with the numbers in the control group.

**Figure 1.**
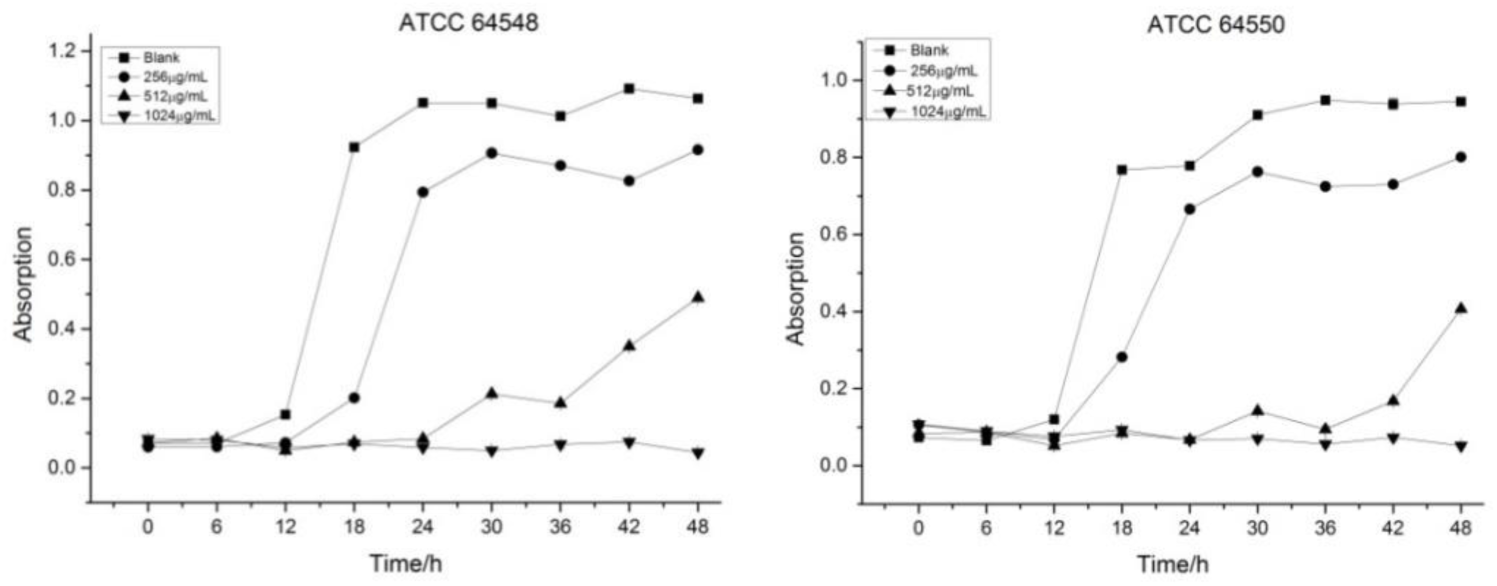
Time-kill curves of ε-PL against *C. albicans* strains ATCC64548 and ATCC64550. Absorption of viable *C. albicans* cells during treatment with ε-PL at 0 (squares), 1/2 MIC (circles), MIC (up triangle), and 2 MIC (down triangle) concentrations in the RPMI-1640 medium (pH 7.0) at 25°C. Error bars indicate the standard deviation.

### ε-PL inhibits *C. albicans* biofilms *in vitro*

The CV method showed that ε-PL eradicated mature biofilms and inhibited biofilm formation (Fig. 2). ε-PL inhibited mature biofilm experiments. First, ε-PL effectively eliminated preformed mature biofilms that had been allowed to grow for 24 h. The effect was dose-dependent (Fig. 2a), and almost all mature biofilms (92%) were removed when treated with 512 μg/mL ε-PL. Second, the biofilm formation test further revealed that the antibiotic membrane effect began at the initial stage of biofilm formation. At 512 and 256 μg/mL concentrations of ε-PL, the biofilm inhibition levels were close to 60% and 73%, respectively (*P* < 0.001). When cells were treated with higher concentrations of ε-PL, no biofilm was formed (Fig. 2b).

**Figure 2.**
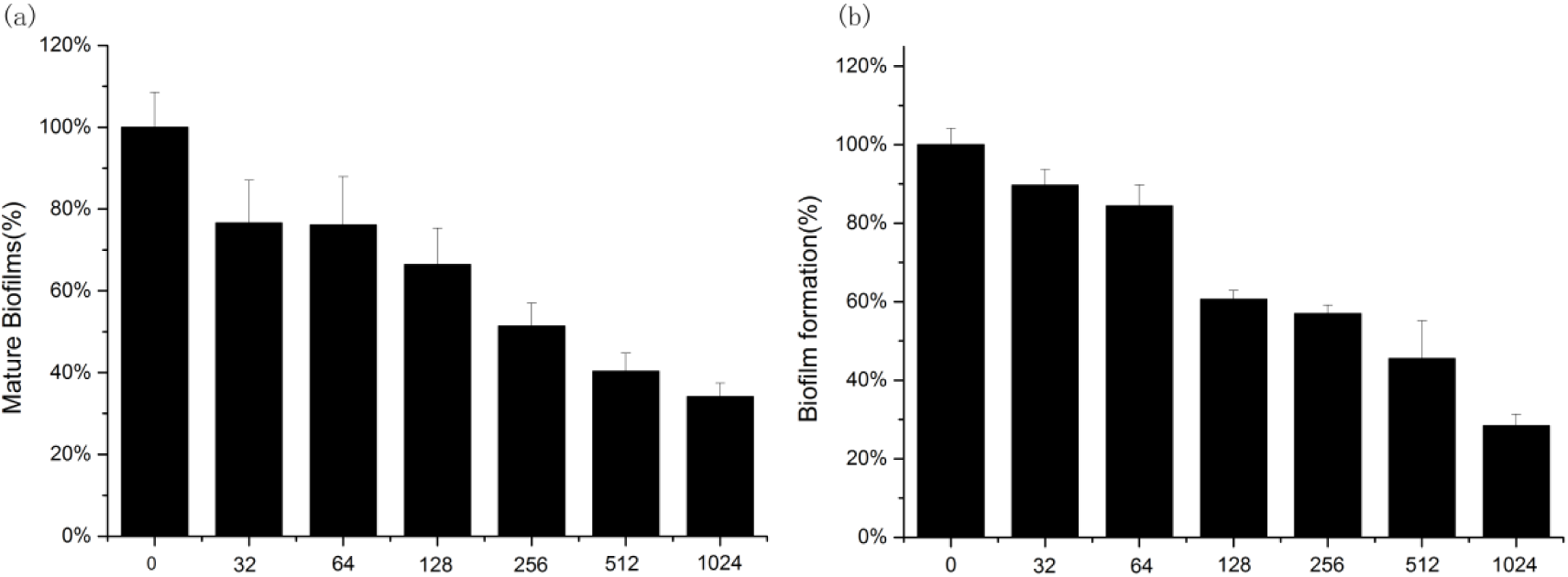
ε-PL inhibited *C. albicans* biofilms *in vitro*. The CV method was performed to determine the effect of ε-PL on mature biofilms (a) and biofilm formation (b). The mature biofilms were left to grow for 24 h, and the biofilm formation assay was conducted after cell adhesion for 1.5 h.

The CLSM analysis further confirmed the anti-biofilm effect of ε-PL. The *C. albicans* biofilm in the control group developed into a crossover network, and the hyphae swelled strongly as shown in Figures 3a and 4a–4c. In the ε-PL treatment group, the degree of biofilm formation damage correlated positively with the drug doses (Figs. 3b–3d and 4d–41). After treatment with 512 μg/mL ε-PL, the activity of hyphae decreased, as visualized using FDA labeling, and the proportion of dead hyphae increased, as analyzed by CLSM (Fig. 3c).

**Figure 3.**
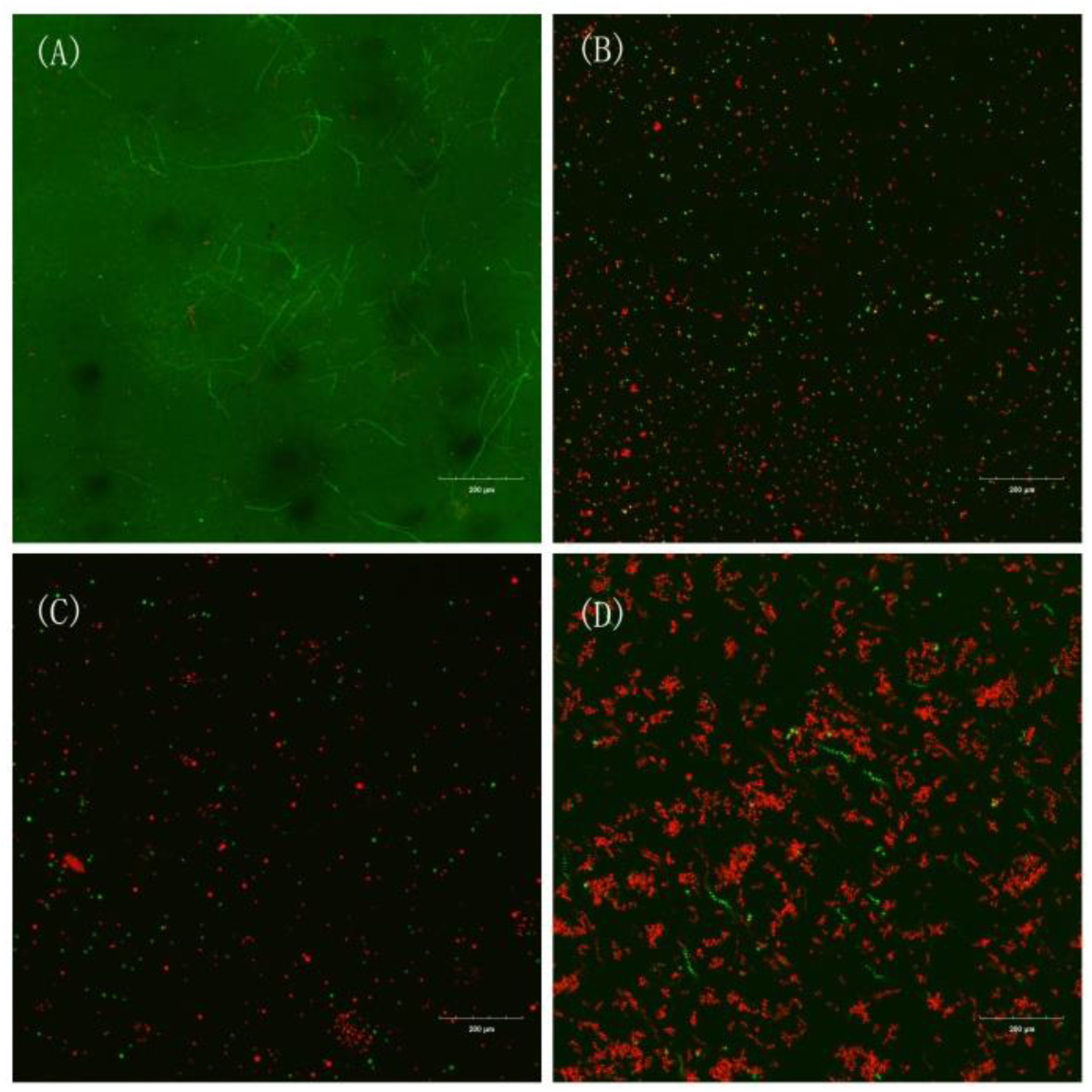
Effects of ε-PL on *C. albicans* biofilm formation shown in CLSM images. (a) Control group; (b) group treated with 256 μg/mL ε-PL; (c) group treated with 512 μg/mL ε-PL; and (d) group treated with 1024 μg/mL ε-PL. FDA and PI were used to distinguish between viable and dead cells.

**Figure 4.**
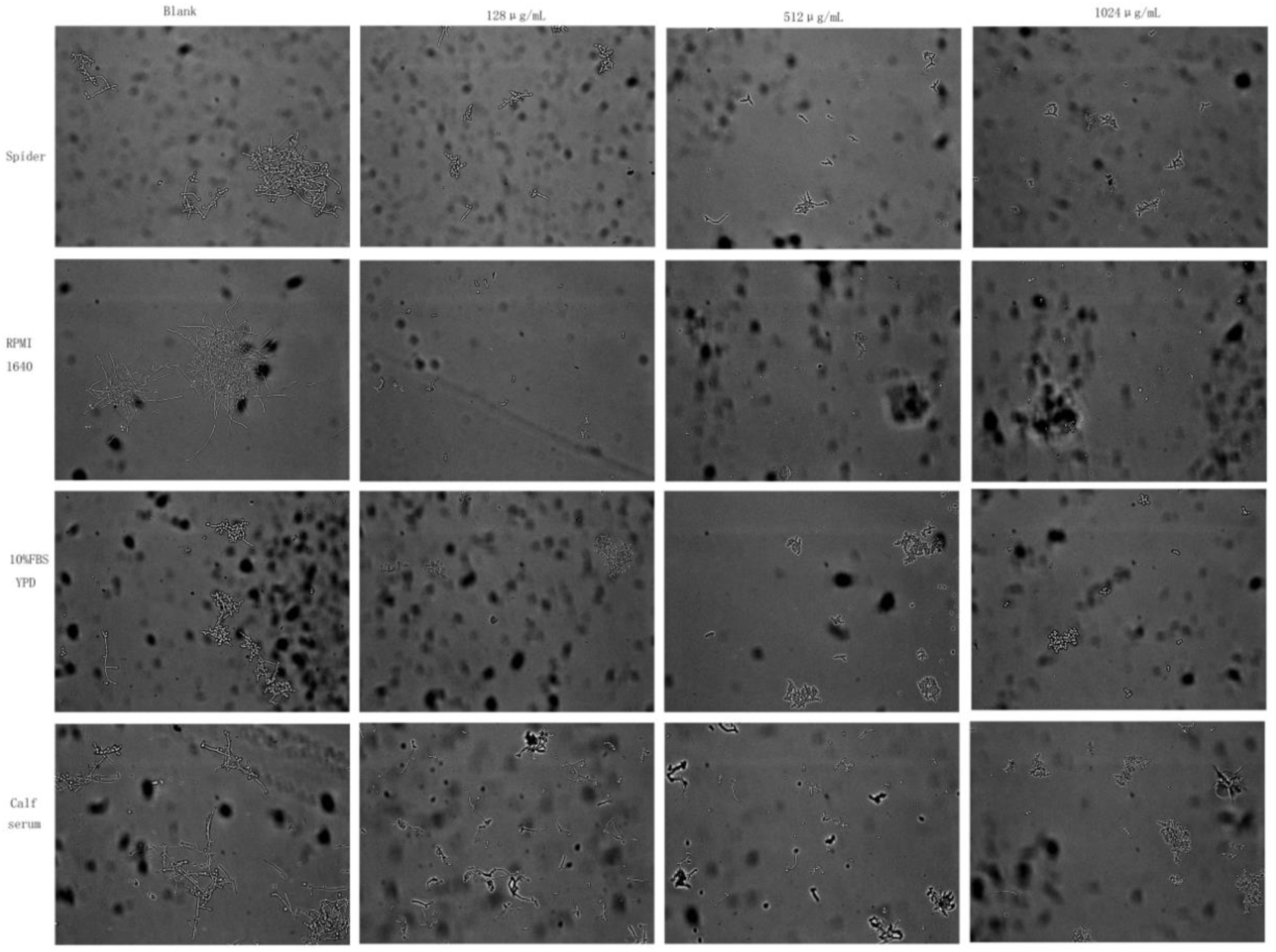
ε-PL inhibited the yeast-to-hypha transition in a dose-dependent manner. Effect of PL on C. albicans hyphal growth. C. albicans cells were grown inbovine serum, YPD (10% serum), RPMI-1640, and Spider media at the indicated concentration of PL at 37 °C for 4 hrs. At the end of incubation an aliquot was withdrawn from each sample and photographed at 400 × magnification.

### ε-PL inhibits the *C. albicans* yeast-to-hypha transition

The hypha of *C. albicans* is an important virulence factor that plays an important role in the pathogenesis. In this experiment, the phase transition of *C. albicans* yeast was observed using an inverted microscope in bovine serum, YPD (10% serum), RPMI-1640, and Spider media for 4 h at 37°C. The results showed that the germination rate and length of *C. albicans* were significantly inhibited by MIC and MFC concentrations, and the inhibitory effect was dose-dependent. The results of the tube showed that the blank group was 4h after the microscope. A large number of hyphae were observed, and the germination rate was low under MIC and MFC concentrations. After 4 h, hyphae appeared in both the low-concentration group and the blank group. MIC and MFC germ tube sprouting rates remained low, and the length of the germ tube was small.

### Effect of ε-PL on oxidative stress of *C. albicans*

The fluorescent probe DCFH-DA was used to measure the content of active oxygen produced by *C. albicans* and different concentrations of technetium-PL after co-culturing at 37°C for 12 h. The relationship between the oxidative damage caused by ROS and the effect of technetium-PL was studied. The nonfluorescent DCFH-DA probe entered the cell membrane freely and was oxidized to DCFH by intracellular lipase. The DCFH probe could not penetrate the cell membrane. It reacted with the active oxygen produced in the cell to produce a fluorescent DCF. The active oxygen content was evaluated by measuring the intensity of fluorescence. Figure 5 shows that the content of ROS produced by different concentrations of ε-PL on *C. albicans* increased with increasing concentrations, compared with the blank group. When the concentration of ε-PL reached MIC, the production rate of ROS accounted for about 93.7%.

**Figure 5.**
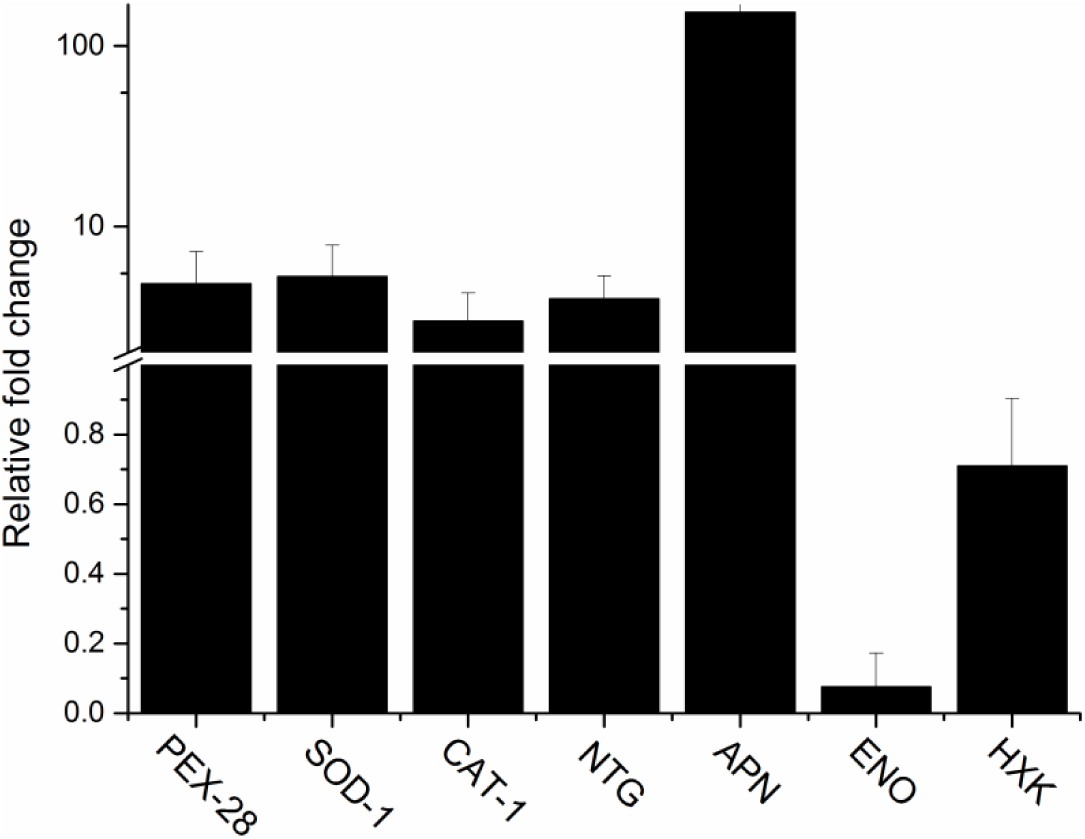
ROS content of *C. albicans* after treatment with different concentrations of ε-PL. Changes in ROS with DCFH-DA staining observed using fluorescence spectrophotometer.

### Lipid oxidative damage to *C. albicans* by ε-PL

To further study the relationship between ROS and membrane oxidative damage of *C. albicans*, the MDA content of *C.albicans* after different concentrations of ε-PL treatment was determined, and the MDA content after adding antioxidant NAC was determined. MDA was the major product of membrane lipid peroxidation and one of the important signs of pressure on *C. albicans* membrane system. By measuring the MDA production, it was safely concluded that different concentrations of ε-PL caused lipid peroxidation of *C. albicans* membranes (Fig. 6). After treatment with 512 μg/mL technetium-PL, the MDA content was about seven times that of the control group. Prior to the addition of antioxidants, the content of MDA positively correlated with the concentration of p-PL. After adding the antioxidant, the MDA content significantly decreased.

**Figure 6.**
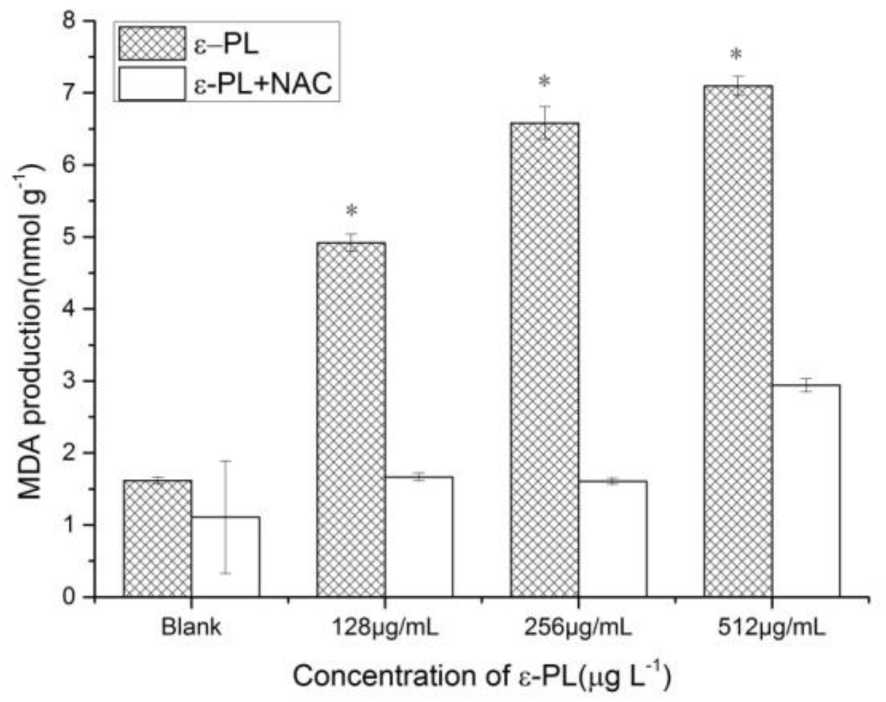
MDA content in *C. albicans* after adding different concentrations of ε-PL (μg/mL).

### Effect of ε-PL on the cell membrane of *C. albicans*

In this study, the effect of ε-PL on the cell membrane integrity of *C. albicans* was determined using the PI method. PI can enter the dead cells and bind to its DNA to form red fluorescence. Measure the ROS reactive oxygen species produced byε-PL induced *C. albicans* by measuring the fluorescence intensity of PI entering the cells.The results showed that compared with the control group, the number of red fluorescent–labeled cells in the ε-PL treatment group increased with the concentration of ε-PL. When the ε-PL concentration reached MFC, the ROS production rate was about 93.7%(FIG 7.0).

**Figure 7.**
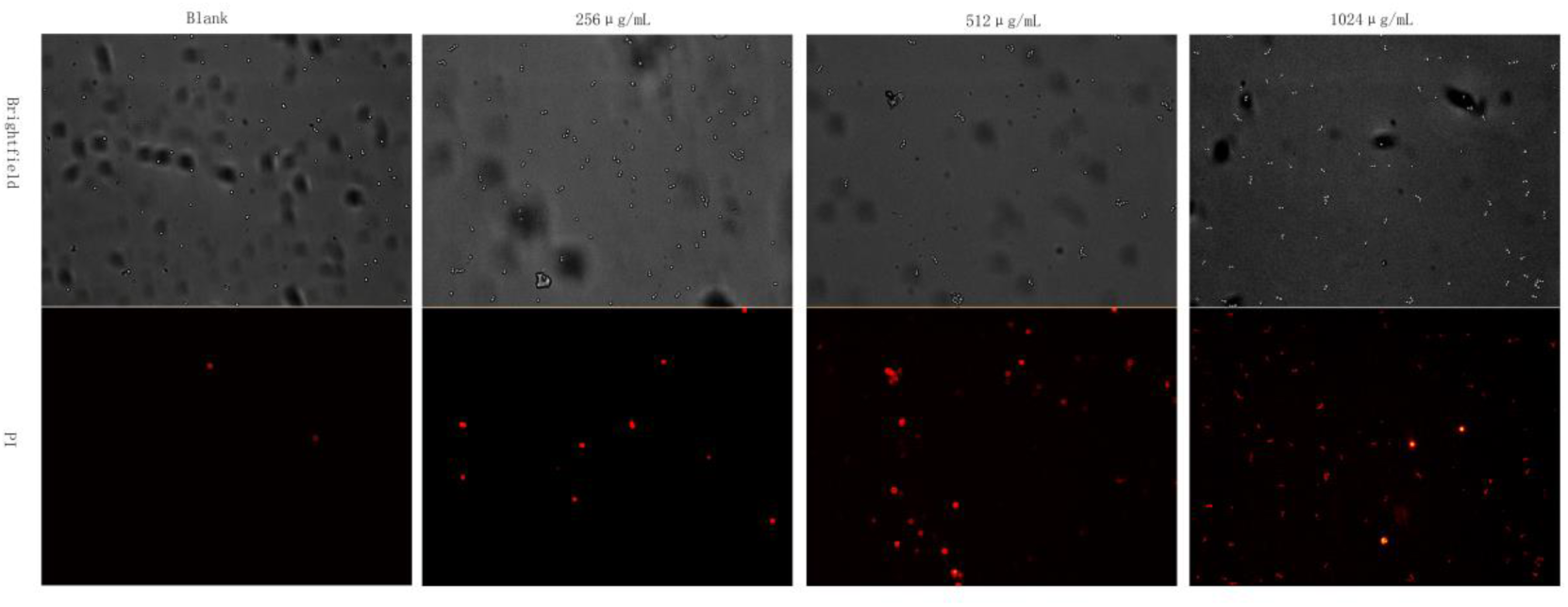
PI percentages of stained *C. albicans* cells after exposure to 0, 256, 512, and 1024 μg/mL ε-PL; the group with no treatments was set as control.

### Effect of ε-PL on the expression of oxidative stress genes in *C. albicans*

The RT-PCR analysis showed that the expression of oxidative stress genes Peroxisomal(*PEX-28*), superoxide dismutase (*SOD-1*), Catalase(*CAT-1*)of *C. albicans* was upregulated by 2.4-fold, 2.6-fold and1.3-fold, respectively, after treatment of *C. albicans* with 512 μg/mL ε-PL. At 1.3 times concentration, the expression of SOS response genes N-glycosylase/AP lyase (*NTG1*) and DNA-(apurinic or apyrimidinic site) lyase(*APN-1*) was upregulated by 1.4-fold and 24.4-fold, respectively, while the expression of *C. albicans* virulence genes enolase(*ENO1*) and hexokinase (*HXK*) was downregulated by 0.09-fold and 0.19-fold, respectively. The results showed that ε-PL-induced damage in *C. albicans* DNA activated the SOS response genes(FIG 8.0).

**Figure 8.**
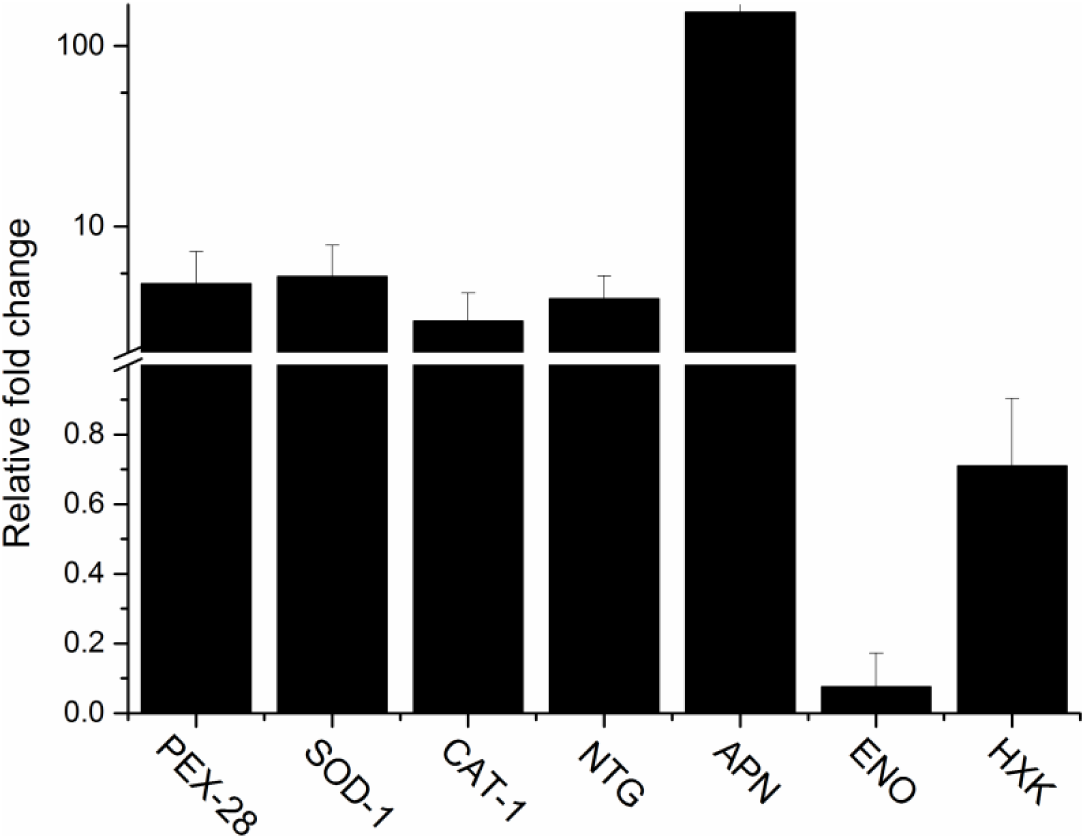
Gene expression levels after ε-PL exposure. RT-PCR was used to validate the differential expression of the *PEX-28, SOD-1, CAT-1, NTG, APN, ENO*, and *HXK* genes. Important genes (*PEX-28, SOD-1*, and *CAT-1*) related to oxidative stress were also quantified by RT-PCR. *NTG* and *APN* genes were closely related to the DNA repair of *C. albicans. ENO* and *HXK* genes were important virulence factors for *C. albicans. ACT-1* was used to normalize the levels of expression.

### Effect of ε-PL on the protein profile of *C. albicans*

The comparative analysis of the bacterial protein profiles of *C. albicans* with different concentrations of ε-PL showed that the characteristic peaks of *C. albicans* mass spectrograms were 6000 *m/z* and 7000 *m/z* after consolidating different concentrations into the same coordinate axis. After treating *C. albicans* with ε-PL, the most obvious change in the protein spectrum was that with the increase in ε-PL concentration, the main peak 6000 *m/z* showed an increasing trend and 7000 *m/z* showed a decreasing trend. The peak value of 4600 *m/z* showed a clear upward trend and simultaneously a significant downward trend at a high concentration. The results of mass spectrometry analysis indicated that ε-PL changed the protein profile of *C. albicans*, while the protein profiles at high and low concentrations were specific(FIG 9.0).

**Figure 9.**
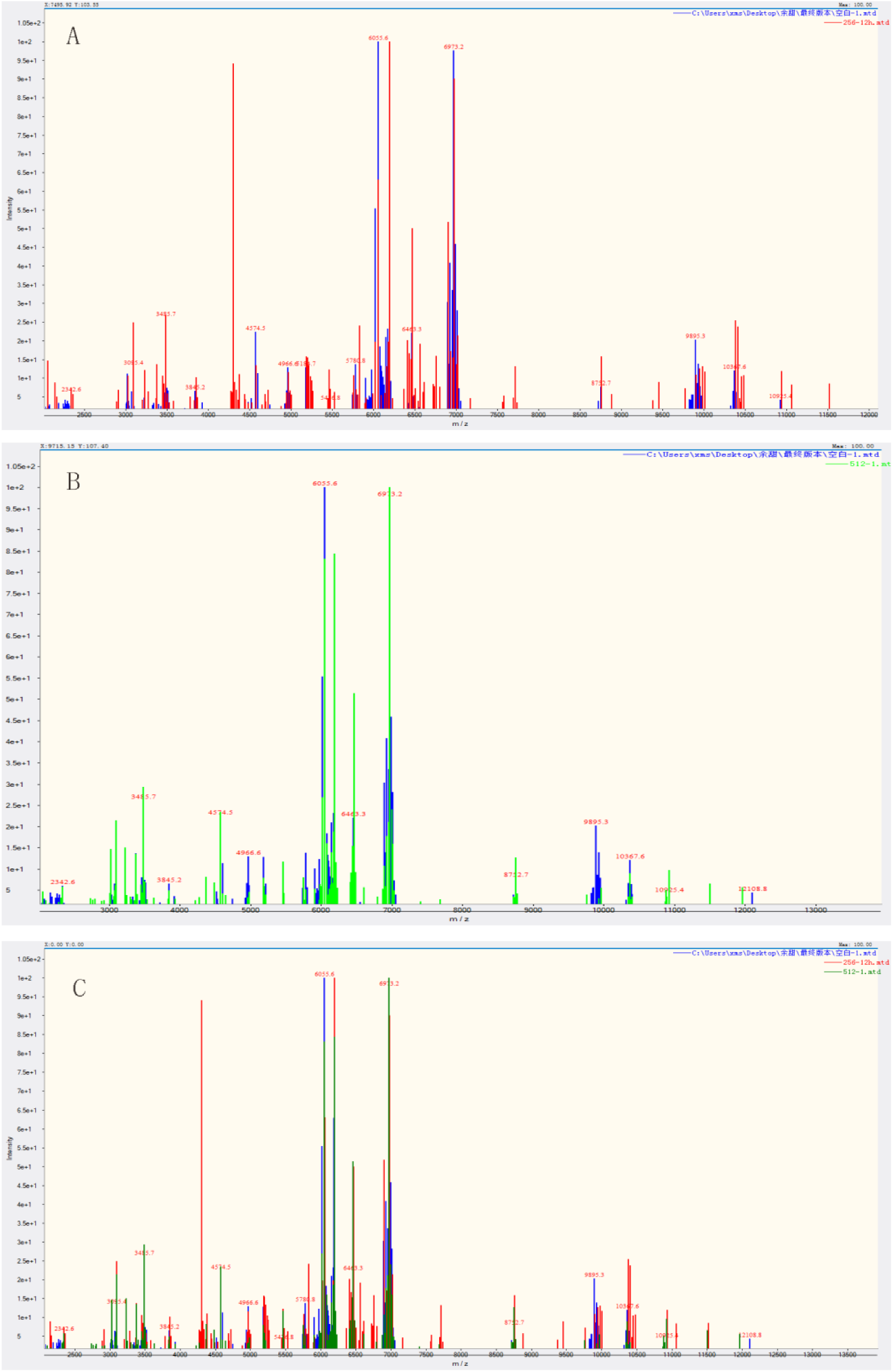
Mass spectrum of *C. albicans* after exposure to different concentrations of ε-PL. Panel A represents the mass spectrum of the blank group (blue) and the 256 μg/mL ε-PL (red) drug treatment group. Panel B represents the mass spectrum of the blank group (blue) and the 512 μg/mL ε-PL (green) drug treatment group. Panel C represents three superimposed maps. This mainly shows the peak changes in the characteristic peaks of *C. albicans* at 6000 *m/z* and 4500 *m/z*.

## Discussion

ε-PL is recognized as a food preservative with broad-spectrum antibacterial activity. However, only the inhibition of fungal growth by a single concentration of ε-PL has been reported so far ^[11]^. This study found that the MIC_80_ and MFC of ε-PL inhibited the growth of planktonic cells of *C. albicans.* Moreover, the time-kill curve indicated that ε-PL inhibited the growth of *C. albicans*, including the fluconazole-sensitive strain ATCC64548 and fluconazole-resistant strain ATCC64550, in a dose-dependent manner. Conazole-resistant strains showed no significant difference. Previous studies showed that biofilm was an important virulence factor of *C. albicans* and is also an important resistance factor ^[21]^. The present study showed that the MIC_80_ of ε-PL significantly prevented the biofilm formation. The SMIC50 assay showed that the MIC of ε-PL significantly inhibited the formation of biofilm, indicating that the inhibitory effect of ε-PL on biofilm was more significant. Zeng ^[22]^ studied the biofilm inhibition of *C. albicans* in sulodexide and showed that SMIC_50_ was usually three to four times of MIC of planktonic bacteria, compared with the inhibitory effect of ε-PL on biofilm in this study.

CLSM and optical microscope analysis of the *C. albicans* biofilm demonstrated that ε-PL inhibited the formation of *C. albicans* biofilm in a dose-dependent manner (Fig. 1). However, the image of MIC_80_ showed the presence of dense hyphae in the absence of ε-PL. The sequential process of biofilm formation was as follows: the adhesion of yeast cells to the substrate, the proliferation of these yeast cells, the formation of biofilms of mycelial cells, the accumulation of extracellular matrix material, and the dispersion of yeast cells from biofilm complexes^[23]^. Hyphae play an important role in the adhesion phase of biofilm formation ^[24]^. Using light microscopy, the yeast-to-hypha transformation of *C. albicans* against *C. albicans* was also compared in the medium that induced mycelial formation, such as bovine serum, RPMI-1640, 10% serum YPD, and Spider media. ε-PL showed a significant inhibitory effect. The hyphae-related experiments clearly demonstrated that ε-PL had a significant inhibitory effect on the yeast–bacteria transformation of *C. albicans*. It was believed that ε-PL reduced the hydrophobicity by inhibiting yeast-to-hypha transformation, preventing biofilm formation, and exerting antibacterial activity^[25]^.

The present study confirmed that ε-PL acted on the cells, and the ROS active oxygen content in *C. albicans* increased. This phenomenon also occurred when ε-PL acted on *E. coli*, which was similar as other antifungal agents reported in the literature. ^[26-27]^. ROS is a class of highly reactive small molecules containing superoxide anion (O^2–^) and hydroxyl radicals (OH^−^)or nonradical molecules such as hydrogen peroxide (H_2_O_2_). The results of this study showed that the contents of ROS and MDA increased with the increase in ε-PL concentration after treating *C. albicans* with ε-PL. After adding the antioxidant, the content of MDA decreased significantly. It was preliminarily confirmed that ε-PL acted on the ROS produced by the bacteria and was closely related to the lipid oxidative damage of the bacterial membrane. In other words, the action of ε-PL on *C. albicans* resulted in excessive ROS production in the bacteria, and the ROS content increased the MDA content in the cell membrane. Bo et al. supported the view that MDA can react with nucleic acids and proteins in *C. albicans* to alter their structural and biological characteristics^[28]^. In this study, PI was used to detect the degree of damage caused by ε-PL on the cell membrane. The results showed that the degree of cell membrane damage of *C. albicans* increased with the increase in ε-PL concentration. It was believed that ε-PL exerted an antibacterial effect on the bacteria mainly by destroying the plasma membrane of the bacteria at high concentrations; at other low concentrations, it acted through other mechanisms^[28]^. In this study, mass spectrometry was used to confirm that the low and high concentrations of ε-PL had an effect on *C. albicans*; mass spectrometry showed significant specificity. Other studies also demonstrated specificity at the genetic level^[29]^.

To further investigate the antibacterial mechanism of ε-PL against *C. albicans*, the expression of oxidative stress–related genes (*PEX-28, SOD-1*, and *CAT-1)* in *C. albicans* induced by ε-PL was examined. *PEX-28* is involved in peroxidase synthesis. It regulates the size and proliferation of peroxisomes^[30]^. *SOD-1* is an important gene of *C. albicans* against oxidative stress^[31]^. The *CAT-1* gene encodes catalase to increase the anti-oxidative stress of *C. albicans*^[32]^. These three important oxidative stress genes showed different degrees of upregulation in this study, indicating that ε-PL exerted oxidative stress on *C. albicans*. The DNA damage repair response (SOS response gene) showed a significant increase in *NTG* and APN genes. The DNA glycosidase encoded by the *NTG* gene and the *APN* endonuclease encoded by *APN* are the major repair genes in the basal excision repair pathway. Repair of DNA damage was caused by endogenous and exogenous ROS^[33]^; the expression of virulence genes *ENO1* and HXK were downregulated in this study. The enolase encoded by *ENO1* gene is considered to be the main antigen in patients with candidiasis. Moreover, it has the advantage of invasion when plasmin binding induces fibrinolysis^[34]^. The *HXK* gene-encoded hexokinase is closely related to the transformation of *C. albicans* to the mycelium form ^[35]^. In conclusion, this study showed that ε-PL could inhibit the ability of *C. albicans* to infect.

## Acknowledgments

This study was supported by the Natural Science Foundation of Gansu Province (17JR5RA035) and the Gansu Provincial Hospital Research Fund (16GSSY1-5). The authors also acknowledge the support extended by the Traditional Chinese Medicine Laboratory of Gansu University and Lanzhou University Laboratory.

